# Causal inference for spatial constancy across whole-body motion

**DOI:** 10.1101/350066

**Authors:** Florian Perdreau, James Cooke, Mathieu Koppen, W. Pieter Medendorp

## Abstract

The brain can estimate the amplitude and direction of self-motion by integrating multiple sources of sensory information, and use this estimate to update object positions in order to provide us with a stable representation of the world. A strategy to improve the precision of the object position estimate would be to integrate this internal estimate and the sensory feedback about the object position based on their reliabilities. Integrating these cues, however, would only be optimal under the assumption that the object has not moved in the world during the intervening body displacement. Therefore, the brain would have to infer whether the internal estimate and the feedback relate to a same external position (stable object), and integrate and/or segregate these cues based on this inference – a process that can be modeled as Bayesian Causal inference. To test this hypothesis, we designed a spatial updating task across passive whole body translation in complete darkness, in which participants (n=11), seated on a vestibular sled, had to remember the world-fixed position of a visual target. Immediately after the translation, a second target (feedback) was briefly flashed around the estimated “updated” target location, and participants had to report the initial target location. We found that the participants’ responses were systematically biased toward the position of the second target position for relatively small but not for large differences between the “updated” and the second target location. This pattern was better captured by a Bayesian causal inference model than by alternative models that would always either integrate or segregate the internally-updated target position and the visual feedback. Our results suggest that the brain implicitly represents the posterior probability that the internally updated estimate and the sensory feedback come from a common cause, and use this probability to weigh the two sources of information in mediating spatial constancy across whole-body motion.

**Author Summary:** A change of an object’s position on our retina can be caused by a change of the object’s location in the world or by a movement of the eye and body. Here, we examine how the brain solves this problem for spatial updating by assessing the probability that the internally-updated location during body motion and observed retinal feedback after the motion stems from the same object location in the world. Guided by Bayesian causal inference model, we demonstrate that participants’ errrors in spatial updating depend nonlinearly on the spatial discrepancy between internally-updated and reafferent visual feedback about the object’s location in the world. We propose that the brain implicitly represents the probability that the internally updated estimate and the sensory feedback come from a common cause, and use this probability to weigh the two sources of information in mediating spatial constancy across whole-body motion.

## Introduction

Motor acts have immediate consequences for the sensory input. For example, a saccadic eye movement across the visual scene temporarily suppresses visual processing [1] and alters the retinal image [2]. Nevertheless, the brain retains correspondence between the presaccadic and postsaccadic scenes – called visual stability – by dissociating these changes in retinal input from those due to changes of the visual scene itself [3].

To do so, it has been suggested that the brain uses an internal forward model that, based on a copy of the saccadic motor command, predicts the postsaccadic scene, which can then be compared with the actual feedback of the postsaccadic scene [4,5]. However, this evaluation process is not flawless because both signals, i.e., the predicted and the actual feedback, are noisy [6]. The optimal strategy for the brain to cope with such uncertainty is through statistically weighting the evidence that the predicted and the actual feedback reflect the same scene or not. This strategy is known as Bayesian causal inference [7].

Recently, we provided evidence for this strategy using the saccadic suppression of displacement task [8], testing how participants judge the presaccadic location of a visual object that shifted during a saccade [9]. Following the rules of Bayesian causal inference, integration was strong when predicted and actual feedback represented spatially close target locations (as if they had a common cause), but weakened with larger spatial differences, depending on the precision of these signals [9].

While the saccadic system has provided evidence for Bayesian causal inference, it is not trivial that this mechanism is also applied to retain visual stability in other motion conditions. Saccades are rapid, self-generated movements that result in an abrupt alteration of the visual scene [10], and more critically in a selective suppression of visual information [11]. Therefore, a mechanism that predicts the reafferent visual information based on motor commands (via forward models) may be a prerequisite for visual updating across saccades [4,5]. In contrast to saccades, passively induced motions, such as riding a car, induce slow and progressive changes of the visual input, and do not have corresponding motor commands that could be used to predict the visual consequences of self-motion. Given these differences, it is not clear how the brain deals with passive self-motion when the environment remains visible.

During passive self-motion, the brain must rely on vestibular and other sensory signals to infer the motion [12–16]. Various studies suggested that there is a clear compensation for passive self-motion in the updating of visual space, although compensation is not always perfect [17–20]. Other studies have shown that this compensation is severely compromised when the vestibular system is lesioned, indicating that vestibular signals weight significantly into visual space updating [21].

Despite these insights, it is important to point out that most of these studies operationalized visual updating by measuring how the brain, in darkness, keeps track of remembered target locations during the motion, in which reliance on self-motion feedback in updating is necessary. In heuristic terms, these self-motion updates may be superfluous in natural settings, where the visual world remains continuously available, uninterrupted by the motion [22].

Here, we ask whether the brain applies Bayesian causal inference in the processing of selfmotion-based visual updates and actual visual feedback signals, or whether it simply derives heuristic, suboptimal solutions to achieve visual stability during passive self-motion, e.g., by relying on visual feedback alone.

To address this question, we designed a spatial updating task across passive whole-body translation, in which participants, seated on a vestibular sled, had to remember the world-fixed position of a visual target and report its location after the intervening body displacement. Critically, in contrast to previous studies, the target was briefly presented again at the end of the displacement (as actual visual feedback), but shifted relative to the updated target location.

In line with the predictions of Bayesian causal inference, we found that our participants’ responses were systematically biased to the actual visual feedback, depending on its spatial discrepancy with the updated location. Our data could not be accounted for by a standard optimal integration model that integrates the internal update and actual feedback irrespectively of their spatial discrepancy, or by reliance on either one of these signals. Our findings suggest that the brain explicitly represents the causal structure in multiple signal integration for visual stability across whole-body motion.

## Methods

### Participants

11 participants took part in the present study (mean age = 27.3 yrs (SE = 2.4), 7 males). All subjects had normal or corrected-to-normal vision and had no known vestibular or neurological disorders. The present study was approved by the Ethics Committee of the Faculty of Social Sciences of the Radboud University, Nijmegen. Every participant gave written informed consent prior to participating in the experiment.

### Apparatus

Participant’s displacement was operated by a custom-made sled, consisting of a chair mounted on an 800-mm track (see [16] for more details). The sled was powered by a linear motor (TB15N, Technotion, Almelo, The Netherlands) and controlled by a Kollmorgen S700 drive (Danaher, Washington, DC). The movements of the sled were controlled with a precision better than 0.034 mm, 2 mm/s and 150 mm/s2. Participants were seated with their interaural axis aligned with the direction of the sled motion. Head movements were restricted by an ear-fixed mold and a chin rest so that participants’ eyes were kept at a distance of 1.47 m orthogonal to an OLED screen of size 1234×676 mm (55EA8809-ZC, LG, Seoul, South Korea). The screen had a refresh rate of 60 Hz. It was placed in front of the sled, aligned with its center (see Fig 1A). A black cardboard frame was mounted on the screen to prevent any residual illumination that could make the screen edges visible.

**Figure 1.**
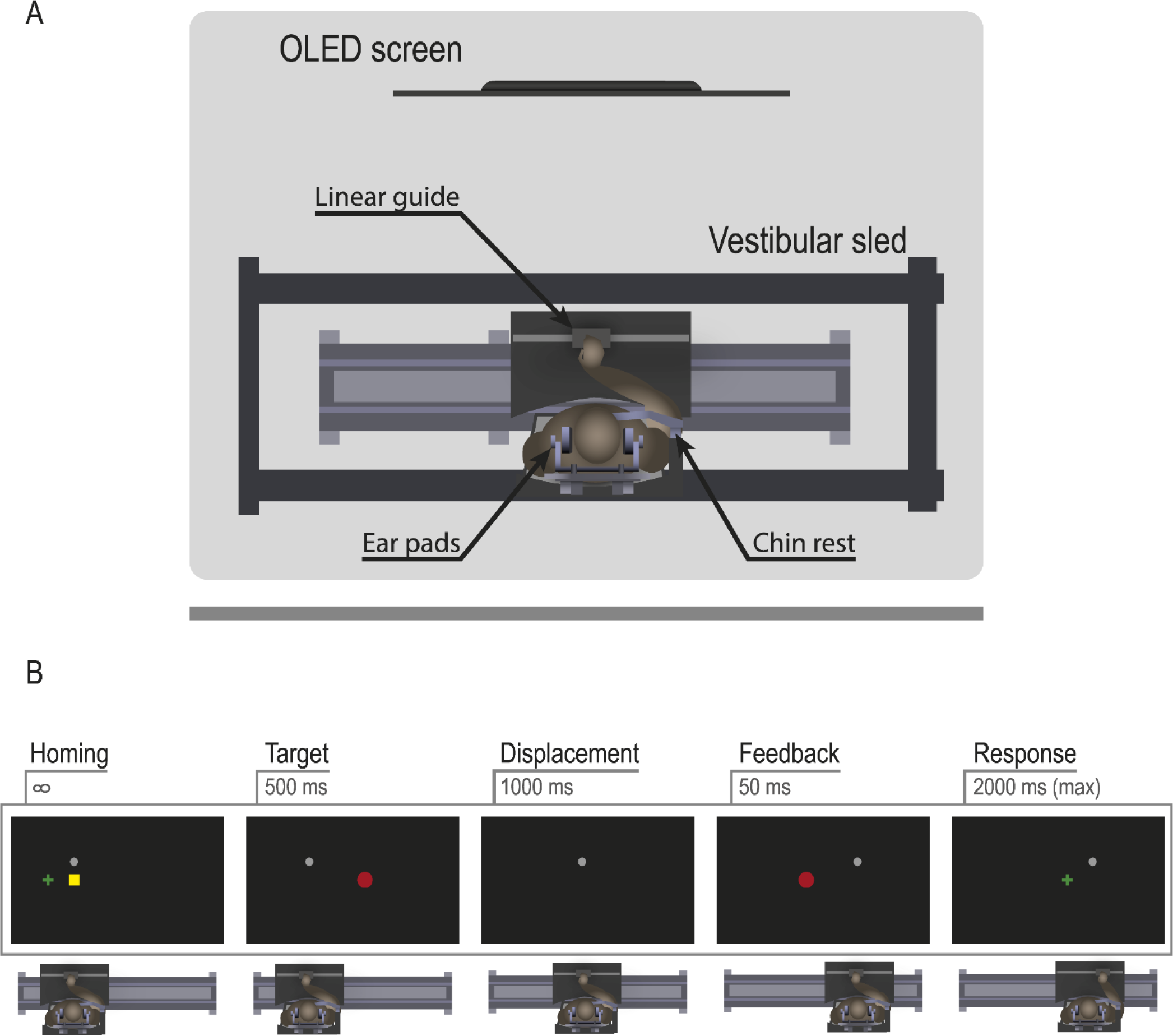
Setup and procedure. (A) Experimental setup. Participants were seated on a vestibular sled. They viewed stimuli on a screen and could control a cursor using a linear guide mounted to the seld. (B) Procedure. A test trial started after the participant had moved a cursor (green cross) to the homing position (yellow square). Next, a target (red disc) was flashed for 300 ms, whose world-centered location had to be remembered. Then, the participant was displaced by 40 cm within 1 s, after which a probe target appeared for 50 ms, shifted relative to the updated target position. Finally, the green cursor reappeared and the participant had to position it at the remembered world-centered location of the target.

A linear guide was mounted on the sled, in front of the participants and at the level of their thoracic diaphragm. By moving a slider on this guide, subjects controled the horizontal motion of a cross-hair cursor displayed on the screen. Position of the slider was continuously tracked at 200 Hz using two Optotrak Certus systems (NDI, Northern Digital Instruments, Waterloo, Canada). The experiment and setup were controlled using software written in Python 2.7.

### Procedure

We designed a task that addresses spatial updating across whole-body passive translation. The task comprised two kinds of trials, *update-only trials* and *test trials.* In the update-only trials, participants had to remember the world-fixed position of a target, briefly flashed prior to the motion, and report its location after the motion. Previous work has shown imperfect spatial updating for passive translation in complete darkness [17,23] and the update-only trials were used to determine the updating gain per participant (see below). The test trials were identical to the update-only trials, with the exception that at the end of the motion, but prior to the participant’s response, the target was briefly displayed again. The location of this probe target varied, but was centered on the internally-updated target location, as estimated on the basis of the update-only trials.

The details of the task are shown in Fig 1B. At the beginning of each trial, participants were passively moved to the homing position of the sled, either at −200 mm or at +200 mm relative to the screen’s center, depending on whether the trial was to test updating across rightward or leftward translation. Then, participants were presented with a 20×20 mm green cursor on the screen, along with a 20×20mm yellow square (cursor homing position) and a gray body-fixed fixation dot (radius: 3.5 mm) that participants had to fixate throughout the trial (Fig 1B). Using the linear guide, participants had to bring the cursor onto the homing position such that both disappeared, which triggered the onset of the target (red disc, radius: 12.5 mm), presented for 300 ms, at one of five possible locations (−100, −50, 0, 50, 100 mm relative to screen center). At target offset, participants were passively moved sideways by 40 cm to the left or to the right for a duration of 1 s with a minimum-jerk velocity profile (peak velocity: 0.7 m/s, peak acceleration: 2.2 m/s^2^). In the test trials, at the end of the motion a probe target was briefly flashed for 50 ms with one of eight possible shifts (−228, −80, −28, −8, 8, 28, 80, 228 mm) relative to the internally updated target location, which was estimated by a preceding block of update- only trials. Finally, in both kinds of trials the cursor reappeared and a brief sound cued the participant to position the cursor at the initially remembered, world-centered location of the target. Participants had 2.5 s to provide their response. If no response was detected within the time limit, the trial was repeated later during the experiment. After the participant had given his/her response, a new trial started, testing updating across motion in the opposite direction. To keep participants motivated and focused on good performance, a message was displayed after every 20 trials showing the average error of the last 20 trials. If the average error was smaller than at the previous message, it was displayed in green (red otherwise). Every participant was instructed to aim for a green feedback signal as often as possible.

The experiment started with a block of 80 update-only trials (eight replications of the five target positions for two movement directions). For each such trial we computed a *motion updating gain* of 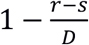, with *s* the actual target position, *r* the response position and *D* the signed motion amplitude *(D* <0 for leftward, *D* >0 for rightward motion) of the trial (all in screen coordinates). This gain equals unity for a perfect motion update and zero if any motion update is lacking. A participant’s updating gain α was determined as his/her average updating gain across all trials of the update-only block and this α was then used to compute his/her internally updated target position in the test trials. Actually, to allow for asymmetric updating, separate updating gains were computed for leftward vs. rightward motion.

Next, the experiment’s *test block* started, consisting of a mix of test trials and update-only trials. Each participant performed 640 test trials (8 replications of 2 motion directions × 5 target locations × 8 probe target shifts). In addition, the test block contained 160 update-only trials, 66% of them randomly interleaved in the first half of the test block and the remaining in the second half. These trials were included to check that the internal update estimate of the preceding update-only block was still valid and to help maintain it [24]. The outcomes of these trials were also used to progressively update the participant’s gain parameter α as the overall average updating gain across all his/her update-only trials.

Target locations and probe target shifts used in the experiment were selected based on pilots and simulations to ensure model recovery.

## Data analysis

### Behavior

For every trial, a response was validated if the cursor reached a velocity of 5 cm/s or less for 300 ms (in screen coordinates). Response error was then computed as the difference between the validated response position and the target position and expressed in mm. Subsequent offline data and statistical analyses were performed using MATLAB (2015b) and R (3.3.2; [25]).

Individual updating gains α were derived as described above and computed separately for leftward and rightward self-motion directions. To determine whether the individual updating gain could be modeled by a single parameter irrespective of self-motion direction we performed a paired samples *t* test on leftward vs. rightward updating gains.

For the test trials, we compared individual response error and variability across conditions by testing linear mixed models using the lme4 R package [26] with motion direction (left, right) and probe target shift (8 levels ranging from −228 to +228 mm) as predictors. In order to compare models, linear and quadratic trends of the probe target effect were investigated. Overall, threshold for statistical significance was defined as 5%.

For the purpose of plotting only, participants’ data were remapped to a rightward body displacement.

### Model

In this study’s main task a trial started with flashing a visual target. Next, the participant was moved sideways and at the end of the motion a probe was flashed either at or with an horizontal deviation from the internally-updated target position (determined using the update-only trials; see Procedure). Throughout the trial a body-fixed fixation cross was present. The participant was then required to indicate the world-fixed position of the target presented before the motion. The purpose was to investigate whether in this situation of passive self-motion and uninterrupted visual input the brain solves the position updating task by combining the available memory and sensory information, on the one hand the internally-updated position of the pre-motion target, denoted **m**, and on the other hand the post-motion probe target position, denoted **ν**, in a statistically optimal fashion, i.e., according to a causal Bayesian inference mechanism [7,9]. The ideas of this approach are now summarized informally, but more details can be found in the supplemental material.

The causal Bayesian inference model is principally probabilistic: both update ***m*** and visual probe percept ***ν*** are considered to be contaminated by noise and represented as probability distributions, taken to be Gaussian. In addition, the model involves a *prior* distribution, also Gaussian, representing the participant’s a priori beliefs about target position, independent of trial information. According to the model, on each trial two hypotheses are considered: one being that ***m*** and ***ν*** have a common cause (here: the probe was displayed at the correct internally updated target position), the other that they have distinct causes (the probe was displaced relative to the internally-updated location).

Under the first hypothesis (***ν*** gives ‘true’ information) the optimal way to combine the ***m, v,*** and prior distributions is by Bayesian integration, resulting in a Gaussian distribution with intermediate mean and higher precision, with precision defined as inverse variance. To be precise, the mean of the integration is the average of the ***m*** and ***ν*** and prior means, each weighted by its own precision, while the integration precision is the sum of the ***m, v,*** and prior precisions. Under the second hypothesis (v is from displaced probe) the optimal way to proceed is simply ignore ***ν*** and just integrate ***m*** and the prior. This is called segregation.

To optimally apply the distributions for the two hypotheses, the integration distribution for a correctly positioned probe, and the segregation distribution for a displaced probe, the probability of the probe being displaced or not is still needed. The model assumes the participant has a prior probability for the probe being correctly positioned, which on each trial is combined with the ***m*** and ***ν*** information of that trial to result in the corresponding posteriori probability. (Qualitatively: the less overlap between ***m*** and ***ν***, the more evidence for displaced ***ν***; again, for the equations see supplemental material). The final model distribution is the mixture of the integration and segregation distributions, each weighted by the posteriori probability of the corresponding hypothesis.

In fitting this model, we have to decide about specifications of the various distributions involved. Priors are regarded as free parameters: the prior probability *p*_*c*_ of correct probe position and the mean π and variance 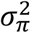 of the prior for pre-movement target position. The distribution ***ν*** for the visually presented probe is supposed to be accurate with variance 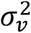 treated as a free parameter.

The distribution ***m*** for the internally-updated position of the pre-motion target cannot be assumed to be accurate: it has been established that, under the conditions of our experiment, passive self-motion amplitude is underestimated [16]. We allow for such underestimation by introducing a gain factor α. For a pre-motion target position ***s*** in body coordinates, the correct update after a movement with amplitude **D** would be ***s*** − **D**. Assuming underestimation of distance **D** by a factor *α* however, the distribution ***m*** is not centered on ***s*** − **D**, but on ***s*** + (1 − α) * *D*. For each participant this gain a was estimated in the first block of update-only trials, which estimate was then used and repeatedly updated in the experiment. The precision of ***m***, on the other hand, contributes another free parameter 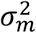.

### Alternative models

The Bayesian causal inference model is a true ‘ideal observer’ model. All available information is used in a statistically optimal way, including computation of the posterior probe displacement probability for determining the weights in the mixture model. Arguably, this theoretical benchmark is hard to attain in real life and the brain might instead resort to some approximating heuristic. One possibility is that it does not compute probabilities for the two competing hypotheses, correct or displaced probe position, but deterministically chooses one or the other, i.e., always integrates or always segregates, respectively. The first alternative, forced fusion, has been previously suggested for saccadic eye-movements [6,27]. It is an extreme case of the causal inference mixture model with 0-1 weights. Similarly, choosing to always segregate corresponds to the opposite 0-1 weighting in the mixture model.

A third possible heuristic is segregation with roles reversed: just process the directly given visual cue and do not consider memory updates. The plausibility of this approach derives from the fact that during passive whole-body movements the brain lacks the possibility to predict the consequence of self-motion due to the absence of motor commands. The usefulness of keeping track of memory and applying updates is not clear for naturalistic situations, where continuous visual feedback is available. For the update-only trials without visual feedback the memory model is retained, based on the assumption that, in the absence of other, more precise information, the brain is able to spatially update an object based on an internal estimate of selfmotion.

Next to the causal inference model also these three simpler models, one integration and two segregation models, were fitted and the model fits were compared.

### Models predictions

Fig 2B shows the predictions of the respective models of the response error (left panel) and response variability (right panel). Generally, the memory-only model (light blue line) corresponds to a flat line (with as intercept the weighted average of one minus the updating gain times motion amplitude and the participant’s prior position, weighted by memory-update and prior precision, respectively). The visual-only model (dark red line) predicts a straight line with slope close to one (visual precision divided by sum of visual and prior precisons). The optimal integration model (orange line) is represented by a straight line with an intermediate slope, determined by the relative precisions of the visual probe target perception, internal memory update, and prior position. The causal inference (green line) prediction is very similar to the integration model close to zero probe target shift (reflecting a high weight for common cause, thus integration), but its slope diminishes in absolute value and may even reverse sign, curving back to the horizontal memory-only axis, reflecting the growing weight for the segregate-memory branch of the mixture distribution with increasing disceprancy of the visual feedback signal.

**Figure 2.**
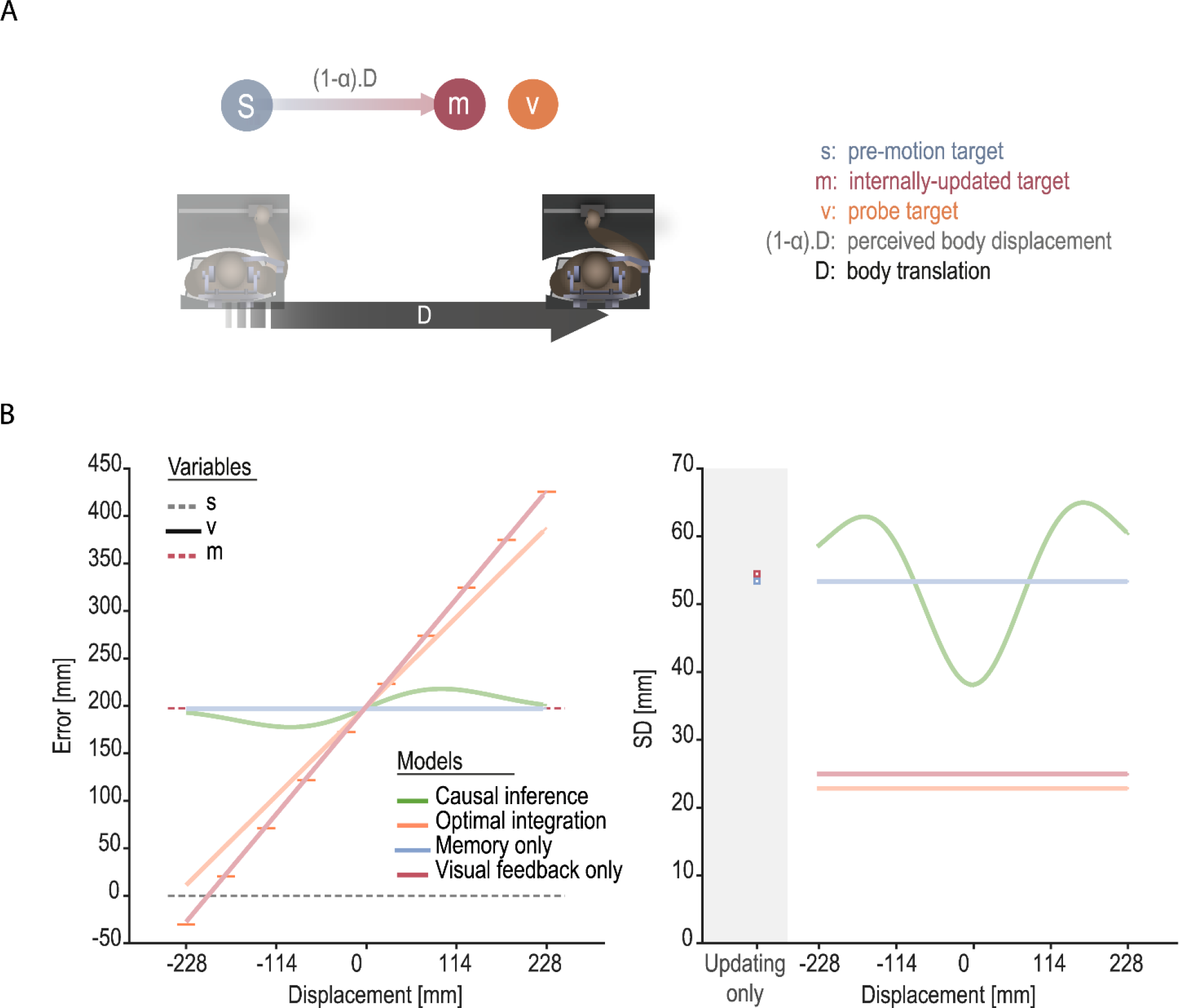
Model’s variables and predictions. (A) Variables: given a target was initially presented at the location ***s***, the task of the observer was to estimate its position after having been passively moved over a distance ***D***. To do so, the observer can use two sources of information: the internally updated target location ***m*** and the probe target position ***v.*** However, because the self-motion is underestimated, the internally updated target location ***m*** would not actually be centered on **s** but rather on *s* + (1 − α) * *D*. (B) Models’ predictions for response error (left) and variability (right): The causal inference model (green) predicts that localization response is biased toward the probe target location (black horizontal lines) for relatively small spatial discrepancies between the probe target (horizontal gray bars) and the internal update *m* (red dashed horizontal line) and becomes aligned with *m* for larger discrepancies. Accordingly, response variability should be greatest for intermediate discrepancies while decreasing as the probe target and the internal update get optimally integrated (small discrepancies) or segregated (large discrepancies). Alternative model are depicted in orange (optimal integration), red (visual feedback only) and light blue (memory-only), see text for further explanation. Because, in these models, *m* and *ν* are always either integrated or segregated according to these alternative models, the corresponding predicted response variability should not depend on the probe shift. Parameters used to generate these predictions are: *σ*_*ν*_ = 25 *mm*, *σ*_π_ = 400 *mm*, *σ*_*m*_ = 60 *mm*, α = 0.5, *p*(*C*) = 0.25, *π* = 0.

Furthermore, the causal inference model also makes testable predictions about response variability (Fig 2B, right panel). The integration branch of this mixture model, which minimizes variability, has a high weight (common cause probability) close to zero probe target shift and this weight decreases with growing spatial discrepancy, i.e., with growing shift amplitude. Therefore, our participants’ response variability should decrease as the spatial discrepancy between the expected and actual feedback decreases. In contrast, the weights predicted by the optimal integration model as well as by the segregation models (memory-only and visual-only) do not depend on the actual spatial discrepancy between the internally-updated target position and the probe position. Therefore, as predicted by these models, response variability as a function of probe displacement follows a flat line whom intercept corresponds depends on the precision of the internal update, of the perception of the probe location and of the prior position (optimal integration), or on the precision of the internal update and of the prior position (memory-only), or on the precision of the perception of the probe location and of the prior position (visual-only).

### Model fitting and evaluation

The causal inference model has six free parameters: the variances 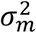 and 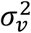 of the ***m*** and ***ν*** distributions, the mean π and variance 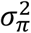 of the target prior, the updating gain α, and the prior probability *p*_*c*_ of correct probe position. The last parameter has no role in the integration model and the visual-only segregation model, for which five parameters are left. The memory-only model has again one parameter less, the variance of the ignored distribution, 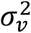, for a total of four parameters. All these models were fit to 1D localization data from both update-only and test trials, which were composed of 8 displacement sizes, 5 targets locations and 2 self-motion directions. Given there is no visual feedback during updating-only trials, we assumed subjects used the memory-model in these conditions to compute their estimate. Thus, the best-fitting parameters set were optimized for both updating-only and test trials.

Parameters were fit by maximum likelihood estimation. Lacking an analytical solution for the mixture model likelihood, this likelihood was sampled by taking 10,000 draws from each of the ***m*** and ***ν*** distributions. Although the segregation and integration processes possess closed-form likelihoods, we used the same simulation approach throughout for consistency. From the discrete draws, a likelihood function was obtained by kernel density estimation (KDE, [28]). We chose a Gaussian kernel with bandwidth *h* determined by Silverman’s rule of thumb (*h* = 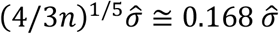, with 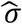 the sample SD; Silverman, 1986). This KDE approach has been used before [29], but alternatively the model draws can be binned into a likelihood histogram [7,9] or the likelihood can be approximated without sampling through numerical integration [30] or linearization [31]. To our knowledge, no study has explicitly compared these approximations.

Likelihood optimization was performed numerically using Bayesian Adaptive Direct Search (BADS, [32]). BADS requires specification of upper and lower bounds as well as plausible upper and lower bounds; the bounds we used can be found in the supplemental material (Table S1). The upper bound for the visual noise parameter (*σ*_*ν*_) was set to 100 mm based on the largest visual noise level estimated in a control condition (see Table S4 in supplemental material). For every model and participant we computed 100 fits using random parameter initializations and selected the best from these. Models’ goodness of fit was assessed using the root mean square error (RMSE). For model comparison the Bayesian Information Criterion (BIC, [33]) was used.

To validate our fitting procedure, we performed both a parameter and model recovery analysis to ensure parameters and models can be inferred well using our experimental design and analysis pipeline (see Tables S2 and S3 in supplemental material).

## Results

The present study aimed at determining the inference mechanism used by the brain to estimate the position of an object after a passive, whole-body translation. More specifically, we were interested in examining whether the brain would only rely on the visual feedback, given it is continuously available during self-motion, or also consider the expected sensory feedback, and then either just use the latter, or always integrate both sources of feedback, or weighting these two possibilities according to the causal inference model. To do so, we designed a spatial updating task across whole-body passive translation where our participants’ task was to remember the world-fixed position of a target displayed prior to the translation and then shown again as a probe at the end of the displacement, at a location shifted relative to the internally updated target position. We were particularly interested in examining the effect of the probe shift on the response bias and variability.

### Estimation of updating gain

For each target position, the internally updated position around which to present the probe target was determined based on the individual updating gain parameter α, during the experiment estimated and updated as the participant’s average across all preceding update-only trials. On average, participants had an updating gain of 0.47 (SE= 0.05). The updating gains between leftward and rightward motion were not significantly different (α_*left*_ = 0.47, α_*right*_ = 0. 46, *t*(10)=0.451, CI(95%)=[−0.018, 0.027], *p*=0.662), so individual updating gains were captured by a single parameter in our models.

### Response error and variability

Fig 3A shows the response error of a typical participant plotted as a function of probe shift from the internally-updated target position (response error of all participants is shown in Fig S2 in supplemental material). While data are replicated from panel to panel, each panel illustrates a specific model prediction. Fig 3B shows the same plots for the group data. While three models predict the relationship to be linear with either a null (memory-only) or positive (visual-only and integration) probe shift slope, causal inference posits a curvilinear relationship with slopes decreasing from zero shift outwards, possibly reversing direction. In a linear mixed model analysis of the response error data, with motion direction and probe target shift as predictors, this opposed curvature (concave down for positive, concave up for negative probe shifts) was modeled by a ‘signed’ quadratic component: the probe shift values squared with a sign reversal added for negative shifts. Motion direction had no main or interaction effect (all *p* > 0.38). Beside a positive linear effect (Χ^2^(1) = 10.23, *p* = 0.001), probe shift also revealed a significant (Χ^2^(1) = 5.37, *p* = 0.020) quadratic effect (with the sign consistent with the CI model). The mixed model required random effects for intercept and both linear and quadratic shift components.

**Figure 3.**
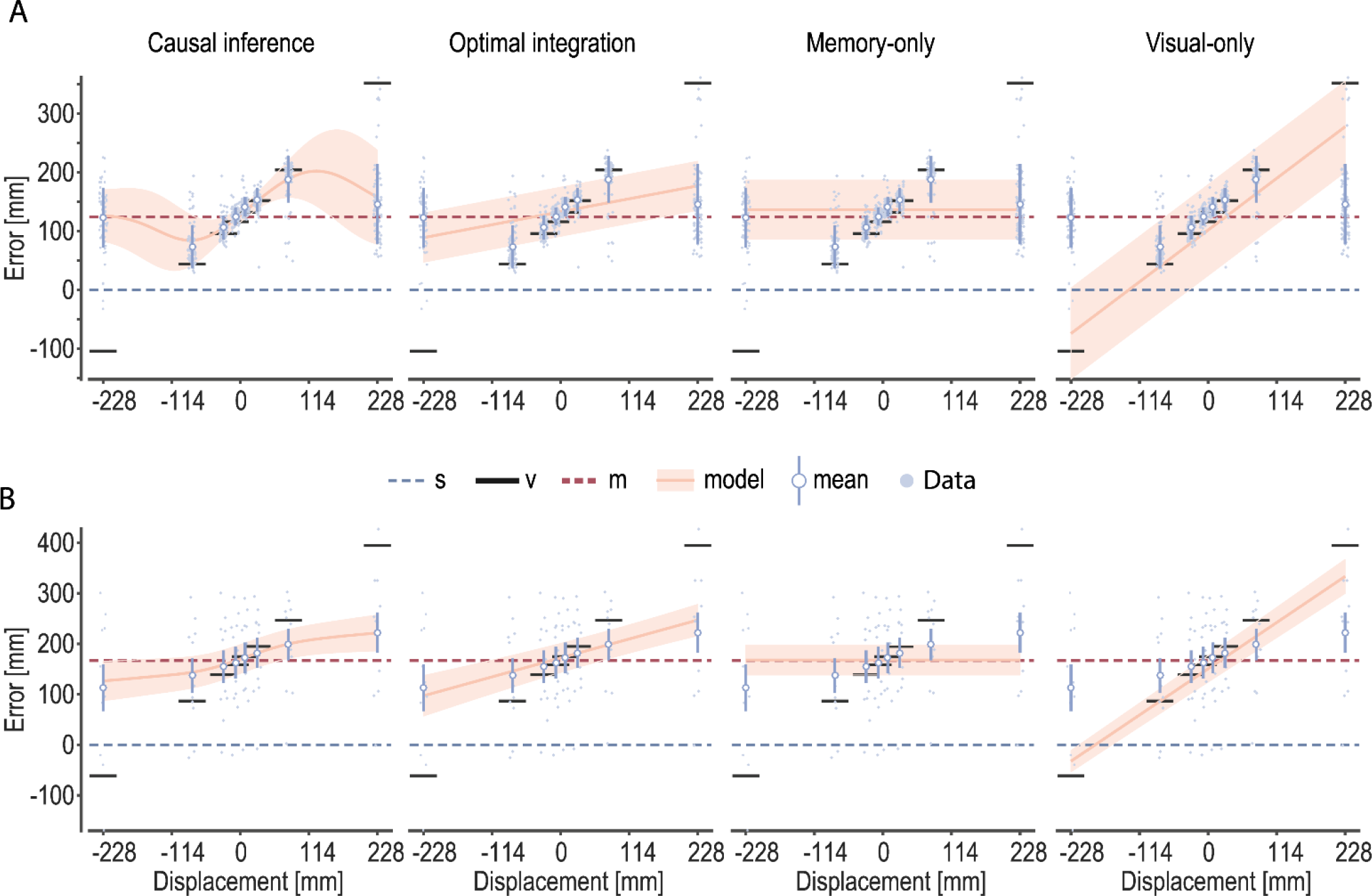
Response error for an individual participant (A) and the group average (B). (A) Response error plotted as a function of the spatial discrepancy between the internally-updated target location and the probe target location together with models’ predictions. Small blue dots represent individual responses, whereas white-filled dots and error bars present the average response and its standard deviation for a specific condition. Solid, orange lines show models’ predictions and colored areas display the standard deviation of the predicted response distribution. Colored dashed line represents the average internally-updated target position, whereas the gray dashed line stands for the target position and the black horizontal bars indicate the probe target position. (B) average participants response error plotted as a function of probe target displacement. Error bars represent the standard error of the mean (S.E.M). Each dot is the average response of one participant. Solid orange line and shaded area represent the average model prediction and the standard error of the mean respectively. Horizontal dashed gray line depicts the target position, whereas the red dashed line and the solid, black lines indicate the average internally-updated target and the probe target positions respectively.

Fig 4A shows the response variability (standard deviation of response error) of an example participant along with our models predictions (one model per panel. Response variability of all participants is shown in Fig S3 in supplemental material). Similarly, Fig 4B presents the same plots at the group level. These variability data were also subjected to a mixed model analysis with motion direction and a linear plus (ordinary) quadratic probe target component as predictors. Again, no main or interaction effect for motion direction was found (all *p* > 0.86). Target shift showed no linear (Χ^2^(1) = 0.24, *p* = 0.621), but a higly significant (x^2^(1) = 62.47, *p* < 10^−14^) concave-up quadratic effect, as predicted by the CI model. Here, the random part consisted of a random intercept only.

**Figure 4.**
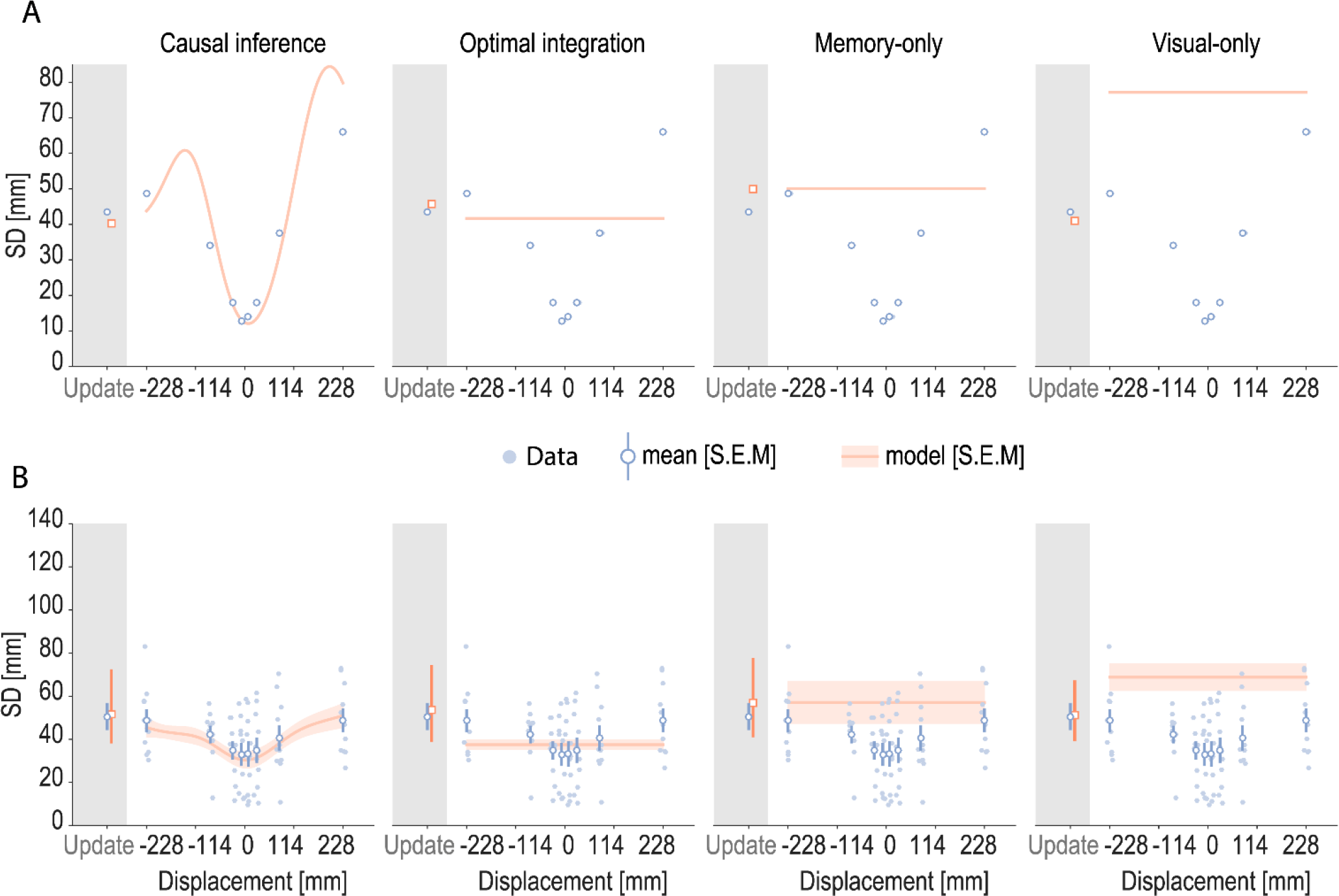
Average response variability for an individual participant (A) and the group average (B). (A) Individual participant’s response variability (standard deviation of response error) and models’ predictions (solid, colored lines) plotted as a function of the spatial discrepancy between the internally-updated target location and the probe target location, as well as for “updating-only” trials (data plotted in gray area). (B) Average participants response variability plotted as a function of target displacement relative to the internally-updated location. The Causal inference model is the only of the tested models that captures the effect of spatial discrepancy on our participants response variability.

### Model fits and evalution

Bayesian causal inference predicts that the memory-based updated target location and the visual feedback are integrated in proportion to the posterior probability that they refer to a same target location (common cause). We tested this hypothesis against the predictions made by three alternative models. One of these is an optimal integration model that combines the memory update and visual feedback based on their respective precisions regardless of their spatial discrepancy. The other two models rely on the heuristic of using just one of both sources: a memory-only model disregarding the probe target and a visual-only model disregarding the internal update. Predictions of the four models are outlined in Fig2B (see Methods).

To quantitatively compare the predictions of our four models at the individual level, we computed model fits per participant. Root-mean-square error (RMSE) computed on each participant’s average response and the model predictions in every condition suggests that our participants’ data were best described by the Causal inference model [RMSE = 12.74 mm, SE = 2.55 mm], followed by the optimal integration model [RMSE = 19.73 (2.54) mm], the memory-only model [RMSE = 37.53 (13.86) mm] and the visual-only model [RMSE = 74.68 (10.00) mm]. A similar analysis applied to our participants’ response variability revealed that the variability pattern (Figs 3B and 4B) clearly could only be captured by the Causal Inference model’s prediction [RMSE= 12.26 (2.72) mm], followed by the optimal integration model [RMSE=12.76 (2.00) mm], the “memory only” model [RMSE=26.94 (10.12) mm], and the visual only model [RMSE=30.91 (SE = 6.59)].

While this fit measure does not take the number of free parameters used by the models into account, we also computed the Bayesian Information criterion (BIC) for each individual model fit. Averaged across participants the causal inference model clearly outperformed the integration model (*ΔBIC* = 294.78, SE = 93.25), the memory-only model (*ΔABIC* = 594.87, SE = 272.06), and the visual-only model (*ΔBIC =*1220.94, SE = 201.87). Given that differences in BIC larger than 20 are considered strong evidence for one model against the other [34,35], this suggests overwhelming evidence for the causal inference model compared to the three alternative models (see Fig 5). Table 1 lists the best fit parameters of the causal inference model and parameter fits of all the models are presented in the supplemental material (Table S5).

**Figure 5.**
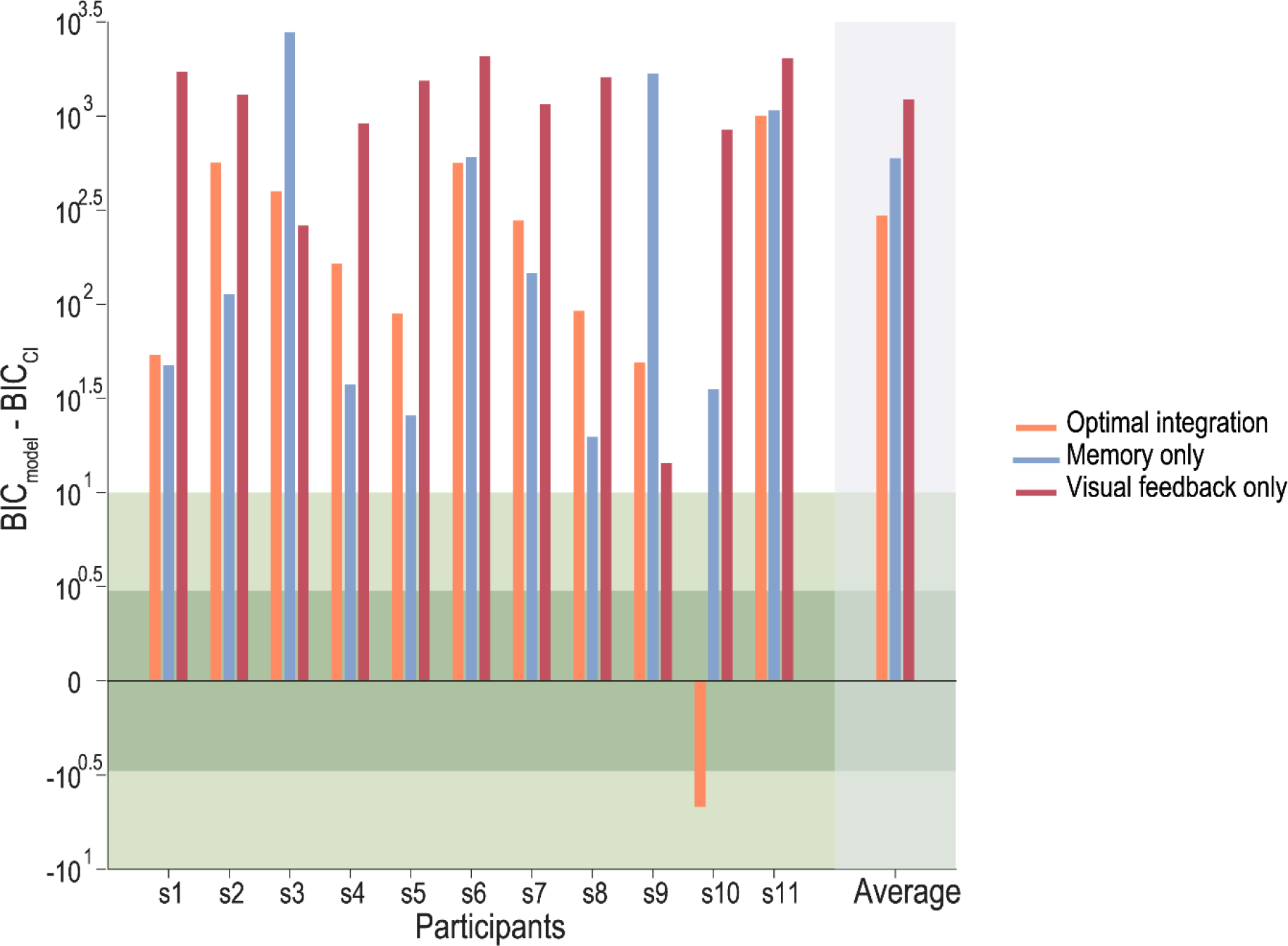
BIC differences between the Bayesian causal inference model and the three alternative models (optimal integration, memory only and visual only). Negative differences would suggest stronger evidence for the alternative model than for the causal inference model and an absolute difference smaller than 10 can be judged uninformative.

**Table 1:** Best parameters recovered by the Causal inference model for every participant.

| *Participant* | $\sigma_v$ (mm) | $\sigma_\pi$ (mm) | $\alpha$ | $\sigma_m$ (mm) | $p(C)$ | $\pi$ (mm) |
| --- | --- | --- | --- | --- | --- | --- |
| *s1* | 25.74 | 364.78 | 0.41 | 32.50 | 0.17 | 93.75 |
| *s2* | 81.26 | 106.94 | -0.02 | 199.98 | 0.06 | -3.41 |
| *s3* | 10.02 | 144.71 | 0.44 | 78.51 | 1.00 | 17.66 |
| *s4* | 28.72 | 128.58 | 1.04 | 61.06 | 0.24 | -22.26 |
| *s5* | 99.90 | 80.31 | 0.05 | 55.66 | 0.68 | 3.08 |
| *s6* | 8.47 | 193.40 | 0.41 | 34.57 | 0.68 | 27.50 |
| *s7* | 3.45 | 170.35 | 0.70 | 72.13 | 0.37 | -25.28 |
| *s8* | 100.00 | 76.61 | -0.08 | 52.40 | 0.38 | 24.97 |
| *s9* | 40.18 | 291.16 | 0.97 | 183.61 | 1.00 | -11.44 |
| *s10* | 100.00 | 76.93 | -0.10 | 77.43 | 1.00 | -3.78 |
| *s11* | 9.92 | 115.81 | 0.62 | 46.86 | 0.89 | -8.67 |
| **Median** | **28.72** | **128.58** | **0.41** | **61.06** | **0.68** | **-3.41** |
| **Mean** | **46.15** | **159.05** | **0.40** | **81.34** | **0.59** | **8.37** |
| **S.E.M** | **12.26** | **28.05** | **0.12** | **17.14** | **0.11** | **10.04** |

## Discussion

Because images from world-centered objects shift on our retina when the eyes move in space, either by eye rotation or body motion, the brain must compensate for our motion in order to keep a spatially constant representation of the world. This spatial updating mechanism involves that the brain internally updates remembered object locations based on the estimated selfmotion. To explain spatial constancy across saccadic eye movements, we recently suggested that the brain relies on a causal inference mechanism [9], showing that the updated target location after saccades and the new visual feedback are integrated and/or segregated based on the posterior probability that they refer to the same position in the world. However, passive self-motion differs from saccadic eye-movements in that visual feedback is typically continously available and spatial updating cannot rely on the efference copy of the motor commands. Therefore, the present study aimed at determining whether the brain would call on Bayesian causal inference or whether it would use a simpler heuristic, for instance solely relying on the available visual feedback for spatial updating after passive whole-body motion. In line with the prediction made by Bayesian causal inference, our results show that responses were biased toward the probe location, with proportionally stronger bias for small probe displacements and a relatively smaller bias for the larger displacements. We conclude that, in order to maintain spatial constancy across passive body motion, the brain would weigh the internally-updated target position and the visual feedback based on the posterior probability that they refer to a common position in the world.

As quantitative support for support this conclusion, we fitted a Bayesian Causal inference model and compared its predictions to those of three alternative models: an optimal integration model and a visual-only as well as a memory-only segregation model. The response patterns of our participants could be better captured by the Causal inference model compared to these alternative models, both in terms of quality of fit and of model evidence as measured by the RMSE and the Bayesian Information Criterion. The fitted prior on the probability of having a common cause was on average about 0.54, but could vary across participants, with some participants being closer to the prediction of an optimal integration model (*p*(*C*) ≈ 1, n = 3) and others being closer to a segregation model (*p*(*C*) ≈ 0, n = 1). Such variability has also been reported in a previous study [9], perhaps related to effects of instruction and experimental context.

Another important prediction of causal inference is that response variability should decrease non-linearly as target and probe are optimally integrated, that is, as its spatial difference of the probe target with the internally-updated target position is smaller. Again, this response variability pattern was observed in most of our participants.

It is important to point out that our data cannot be used to determine whether our participants were using a true Bayes-optimal causal inference strategy or a suboptimal approximation of it
[36]. More specifically, because the posterior probability of having a common cause partially depends on how these two cues can be spatiotemporally discriminated, this would depend on how precise their representation is. Our previous study investigating the role of causal inference in spatial constancy across saccades manipulated the sensory reliability of the post-saccadic probe target by varying its presentation duration [9]. It was found that for longer presentation durations, which results in a more precise probe target representation, participants’ responses were more biased toward the probe location for small displacement (stronger integration) but also more quickly disregarded as the size of the displacement increased. This result suggest that observers could obey a true Bayes-optimal decision strategy in spatial constancy tasks.

While our results suggest updating responses rely on a Bayesian causal inference, they cannot disentangle the different possible implementations of the decision strategy used in causal inference. Here, we only considered causal inference as a model averaging approach: both integration and segregation estimates are combined and weighted based on the posterior probability of having a common cause in a particular trial. Alternative decision strategies have been suggested as well and have been compared in previous studies, including model selection [37], probability matching [9,38] and coupling prior [39,40]. Nonetheless, model averaging most of the time outperformed these alternative decision strategies in predicting participants response in cue-combination and unity judgment tasks [7,9,30,37].

Previous research has investigated spatial updating across body translation in complete darkness using psychophysical procedure where participants have to compare the position of a probe target to a reference [17,19]. Despite the similarity with our test trials where a probe target was shown at the end of the translation, the aim of these studies was in fact closer to the goal of our updating-only trials: measuring how an observer updates an object position based on vestibular cues only. A critical difference with our test trials lies in the role played by the probe target and the way it can be used. Comparing the probe position to the target position necessarily implies to segregate these positions and to regard them as being generated by different causes (compare one object vs. another). In contrast, we used an estimation task where participants had to report the location of the target, possibly combining the probe location as an additional source of information. In contrast to using a 2-AFC task as previously described, our estimation procedure allows us to determine the way visual feedback regarding the target can be used in order to more precisely update the target location.

Our spatial updating task involved passive linear body-translation in complete darkness with restricted eye-movements due to the body-fixed fixation target. This allowed us to better control the types of sources of information that the brain could use in order to estimate the translation amplitude and consequently update the target location. Optic flow was not available at all. Therefore, the amplitude estimate could mostly be derived from the integration of canals and otoliths signals about the angular and linear acceleration of the head in space respectively [15,41]. Increasing the number of sensory and motor sources about self-motion, reflecting more ecological situations, however, would not rule out the use of causal inference. Integration of these multiple sources of information would actually result in a more precise estimate of the updated target location, which would then be better discriminated from the sensory feedback of the target. The sharpening of the sensory cue estimates should finally impact on the posterior probability of having a common cause and, consequently, on the weighting of integration and/or segregation of these estimates. Interestingly, this causal inference could take place at multiple stages of processing: e.g. related to the multisensory cue-combination to estimate self-motion, and related to the integration of the sensory feedback in order to improve the estimate of the target location or to detect an external change in the object.

We used a computational approach to describe and predict human updating behavior. Previous theoretical work suggested that Bayesian optimal integration could be supported by the linear combination of neural population activity that can be approximated by Poisson-like distributions [42,43]. In line with this suggestion, it has been shown in macaques that MSTd neurons compute the weighted sum of their inputs, the weights of which were varying according to motion coherence, which was a manipulation of cue reliability [44]. Similar evidence for neuronal correlates of optimal integration has been found in studies involving visuo-vestibular stimuli [45]. More recently, an audio-visual cue-combination study combined with fMRI found evidence for a possible cortical hierarchy implementing causal inference [46]. At the bottom of the hierarchy, primary sensory areas encode their preferred stimulus (segregation estimates), whereas optimal integration of the sensory cues and the encoding of the uncertainty about their causal structure would occur higher in the hierarchy (posterior intraparietal sulcus and anterior intraparietal sulcus respectively). This study, however, considered Bayesian causal inference in a task that involves the combination of unisensory cues. In contrast, spatial updating involves the combination of a sensory feedback and an internal, amodal estimate of the expected sensory feedback itself derived from multisensory sources of information. Therefore, it remains to be determined whether this cortical hierarchy would also support Bayesian causal inference for spatial constancy. Interfering techniques, such as TMS, could be used in order to test whether it affects how participants integrate sensory cues given their spatial disparity in spatial updating tasks.

## Acknowledgments

This research was supported by a VICI grant to W.P.M.

## Author contributions

Conceived and designed the experiments: FP JC WPM. Performed the experiments: FP. Analyzed the data: FP JC MK WPM. Wrote the paper: FP JC MK WPM.

